# Continuous Evaluation of Ligand Protein Predictions: A Weekly Community Challenge for Drug Docking

**DOI:** 10.1101/469940

**Authors:** Jeffrey R. Wagner, Christopher P. Churas, Shuai Liu, Robert V. Swift, Michael Chiu, Chenghua Shao, Victoria A. Feher, Stephen K. Burley, Michael K. Gilson, Rommie E. Amaro

## Abstract

Docking calculations can be used to accelerate drug discovery by providing predictions of the poses of candidate ligands bound to a targeted protein. However, studies in the literature use varied docking methods, and it is not clear which work best, either in general or for specific protein targets. In addition, a complete docking calculation requires components beyond the docking algorithm itself, such as preparation of the protein and ligand for calculations, and it is difficult to isolate which aspects of a method are most in need of improvement. To address such issues, we have developed the Continuous Evaluation of Ligand Protein Predictions (CELPP), a weekly blinded challenge for automated docking workflows. Participants in CELPP create a workflow to predict protein-ligand binding poses, which is then tasked with predicting 10-100 new (never before released) protein-ligand crystal structures each week. CELPP evaluates the accuracy of each workflow’s predictions and posts the scores online. CELPP is a new cyberinfrastructure resource to identify the strengths and weaknesses of current approaches, help map docking problems to the algorithms most likely to overcome them, and illuminate areas of unmet need in structure-guided drug design.

## 2 Introduction

The discovery of a small molecule that binds a disease-related protein with high affinity is a key step in many drug discovery projects. In the pharmaceutical industry, this step has been estimated to require over three years of work on average, at a net cost per launched drug rivaling that of clinical trials.^1^ The process is perhaps most efficient when a high-resolution structure of the targeted protein is available, such as from X-ray crystallographic studies. In this setting, structure-based computational methods may be used to accelerate the discovery of high affinity ligands.^2–13^ The computational challenge of structure-based ligand design may be considered to comprise two main components. The first is prediction of the bound conformation, or pose, of a candidate ligand, typically by fast, ligand-protein docking algorithms.^14–28^ The second involves using the predicted pose to assess the candidate ligand’s binding affinity for the targeted protein, or at least to rank its affinity relative to the affinities of other compounds one contemplates purchasing or synthesizing. Both components have been the subject of intensive research and development in both academic and commercial settings.^29–44^ Nonetheless, computational methods for pose prediction and affinity ranking have yet to fulfill their perceived promise, as neither is yet fully reliable.^45–51^ In fact, it is surprisingly difficult even to compare the reliability of various methods in a consistent manner, and this limitation makes it correspondingly difficult to make and verify technical progress. Part of the challenge of rigorously comparing methods relates to reproducibility (or lack thereof) of the complicated and highly variable end-to-end computational experiments required for pose and affinity prediction.^52^ To address these issues, the community has seen a dramatic uptick in the use and availability of automated workflows that clearly memorialize a particular experiment and provide different approaches for their execution and deployment.^53–57^

Further, although many performance comparisons have been published, the results can be difficult to interpret.^47, 58–68^ For example, new docking algorithms are frequently published along with a comparison against existing methods, but this comparison is often secondary to the description of the new algorithm, and hence not fully developed. Additionally, different methods are typically tested against different sets of protein-ligand complexes, so a consistent set of comparisons may not be available. Finally, even when a study carries out careful benchmarking of multiple methods against a common dataset, the dataset often contains protein-ligand cocrystal structures that have already been published. Such retrospective studies are suboptimal, because they risk unintentional bias and because structures in the test set might have been used previously in training the docking algorithms.^69^

Several initiatives have addressed these limitations through prospective, or blinded, prediction challenges. In such challenges, researchers evaluate methods against a common set of test cases for which the experimental structures are withheld until after the computational predictions have been made. Prior blinded challenges include the GSK challenge,^47^ CSAR,^66, 70–73^ and GPCRDOCK.^74–76^ Similarly, in recent years, the Drug Design Resource (D3R) has run blinded prediction challenges called the Grand Challenges.^45, 46^ These efforts have led to useful benchmarking strategies, provided insight about best practices, and sometimes yielded unexpected results regarding the effectiveness of various technical approaches. However, such episodic challenges have not been large and systematic enough to afford statistically meaningful distinctions among individual methods or to support an efficient cycle of development and evaluation that can persistently accelerate progress in the field.

Here, we introduce a new blinded prediction challenge which overcomes these limitations by taking advantage of ongoing available data streams and advances in data science.^77–79^ The Continuous Evaluation of Ligand Protein Predictions (CELPP) challenge uses the Protein Data Bank’s (PDB)^80–84^ weekly publication of a list of structures slated for imminent release into the public domain as the basis for a weekly, pose-prediction challenge. This rolling challenge is akin to the Continuous Automated Model Evaluation (CAMEO) protein structure prediction challenge,^85^ which served as its inspiration. The following sections detail the structure of the CELPP challenge, the automation used to enable smooth weekly operations, initial results for a number of docking workflows, and implications and directions for this community science project.

## 3 Methods

### 3.1 Overview of the CELPP Blinded Cross-Docking Challenge

Each week (Figure 1), in-house CELPP scripts download the list of new PDB entries to be released five days later and identify those which contain protein-small molecule cocrystal structures suitable for automated docking calculations (https://github.com/drugdata/D3R). At this stage, the only information available about each of these structures, called target complexes, is the identity of the ligand, the amino acid sequence of the protein, and the pH of the mother liquor from the crystallographic study. Additional scripts then search existing PDB entries for crystal structures of each target protein and extract up to five structures appropriate for docking calculations, as detailed in Section 3.2.1. These protein structures are incorporated into the weekly CELPP data package, along with the ligand identities, crystallization pH values, and additional information (Section 3.2.2). CELPP participants download the data package, run their own workflows to predict the ligand binding poses, and submit their predictions to a personal, password-protected web directory before the deadline; i.e., shortly before release of the new PDB entries containing the actual crystallographic poses. Following the deadline, D3R scripts evaluate the submitted predictions, send the evaluation results to each participant, and add the results to running statistics available online (http://drugdesigndata.org/about/celpp).

**Figure 1.**
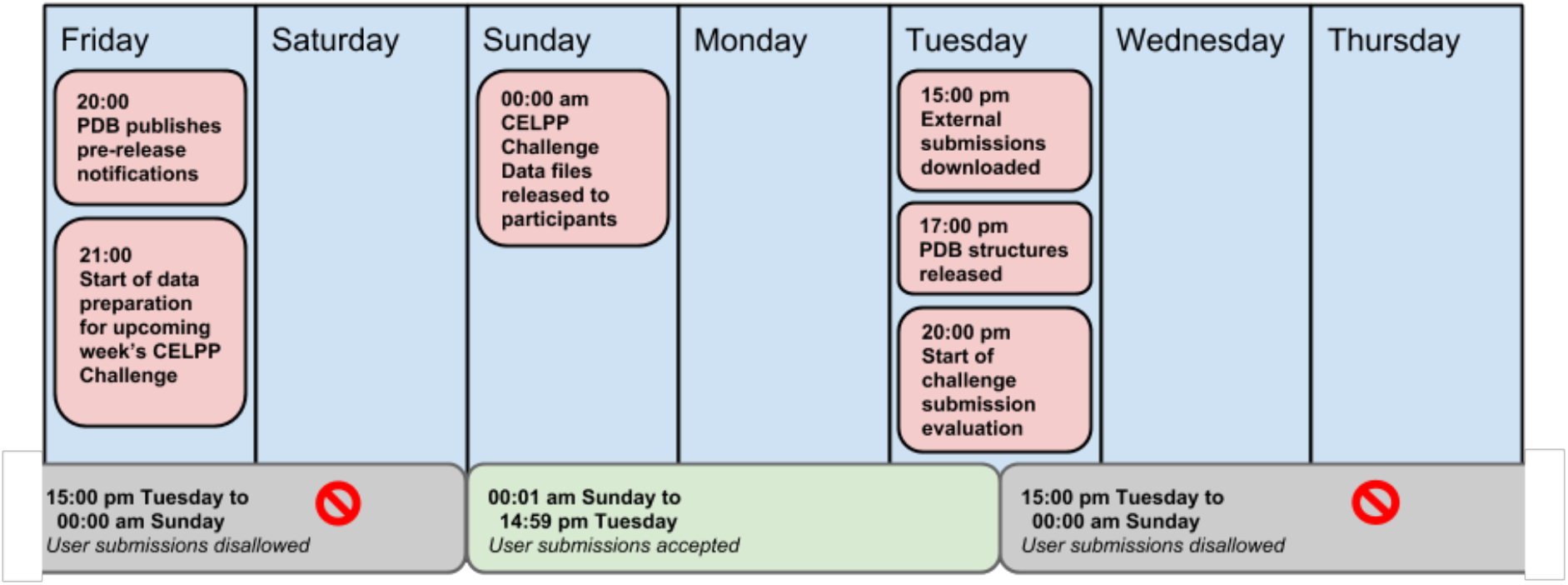
The CELPP week. The CELPP week begins with the publication of PDB pre-release data on Friday evening. Challenge data preparation runs Friday evening and Saturday, and the upcoming week’s challenge package is made available to participants by the beginning of Sunday. Submissions are then accepted until Tuesday at 3:01 pm. Evaluation of the predictions begins on Tuesday evening, following release of the new PDB entries used in the challenge. Times are in the Pacific time zone.

It should be evident from this description that CELPP presents what is known as a *cross-docking* challenge, i.e., the protein structure into which the ligand is docked was previously determined either with a different ligand or with no ligand at all.^17, 86, 87^ This may be contrasted with the *self-docking* problem, in which the ligand is docked back into the protein structure resolved in complex with the same ligand. Cross-docking is typically more difficult, because the protein structure used has not adapted to the ligand being docked, and indeed may be in a conformation that the ligand does not fit well. However, cross-docking is arguably a more important problem than self-docking, because it models the real-world applications of docking methods, where the entire purpose of docking is to avoid having to determine cocrystal structures for every ligand of interest in a drug discovery project.

The success of a docking calculation depends not only on the algorithm itself, but also on other methods and parameters in the overall workflow. For example, in cross-docking, one of the most important decisions is what existing structure of the protein to use in the calculation.^45, 46^ Additional issues arise in the preparation of the protein and ligand structures for docking. For the protein, it is often necessary to decide how to resolve ambiguously assigned electron density, whether to remove or retain specific solvent molecules, how to account for crystallization artifacts (such as non-natural solvents, crystal contacts, and non-physiologic temperature), whether to select alternate side-chain conformations, and how to set the protonation states of titratable residues.^62, 88–91^ For the ligand, issues may include assignment of protonation and tautomer states, and the conformations of flexible rings.^88, 92, 93^

### 3.2 Hosting the Challenge

#### 3.2.1 Selection of target complexes and receptor structures

Every Friday, the PDB provides files (http://www.wwpdb.org/files/) listing the new crystal structures that will be released the at the end of the following Tuesday (Figure 1). For each forthcoming PDB entry, this pre-release notification contains the PDB ID, the protein sequence (s), the identities of the ligands, if any, in the form of InChi strings,^94^ and the pH at which the structure was determined. To be designated as a CELPP “target”, a structure must include a single druglike ligand (see below). There also must be at least one X-ray crystal structure of the same protein extant in the PDB to serve as a suitable “candidate” for cross-docking, and the target must have only one unique protein sequence, to avoid situations in which the target and candidate ligands are bound to different binding sites. (See Figure 2.) A ligand is considered drug-like if it is not a single metal ion or typical solvent molecule, and if it is not on an exclusion list of common cosolutes and cofactors (e.g., Zn^++^, ethylene glycol, and NADH; see Scheme S3). We also exclude ligands with more than 100 self-symmetries (i.e., automorphs), because evaluating symmetry-corrected root-mean-square deviations between predicted and crystal poses becomes excessively time-consuming in such cases.^95^ Finally, we include one “standard” target, PDB ID 1FCZ, in the challenge package each week to monitor workflow stability.

**Figure 2.**
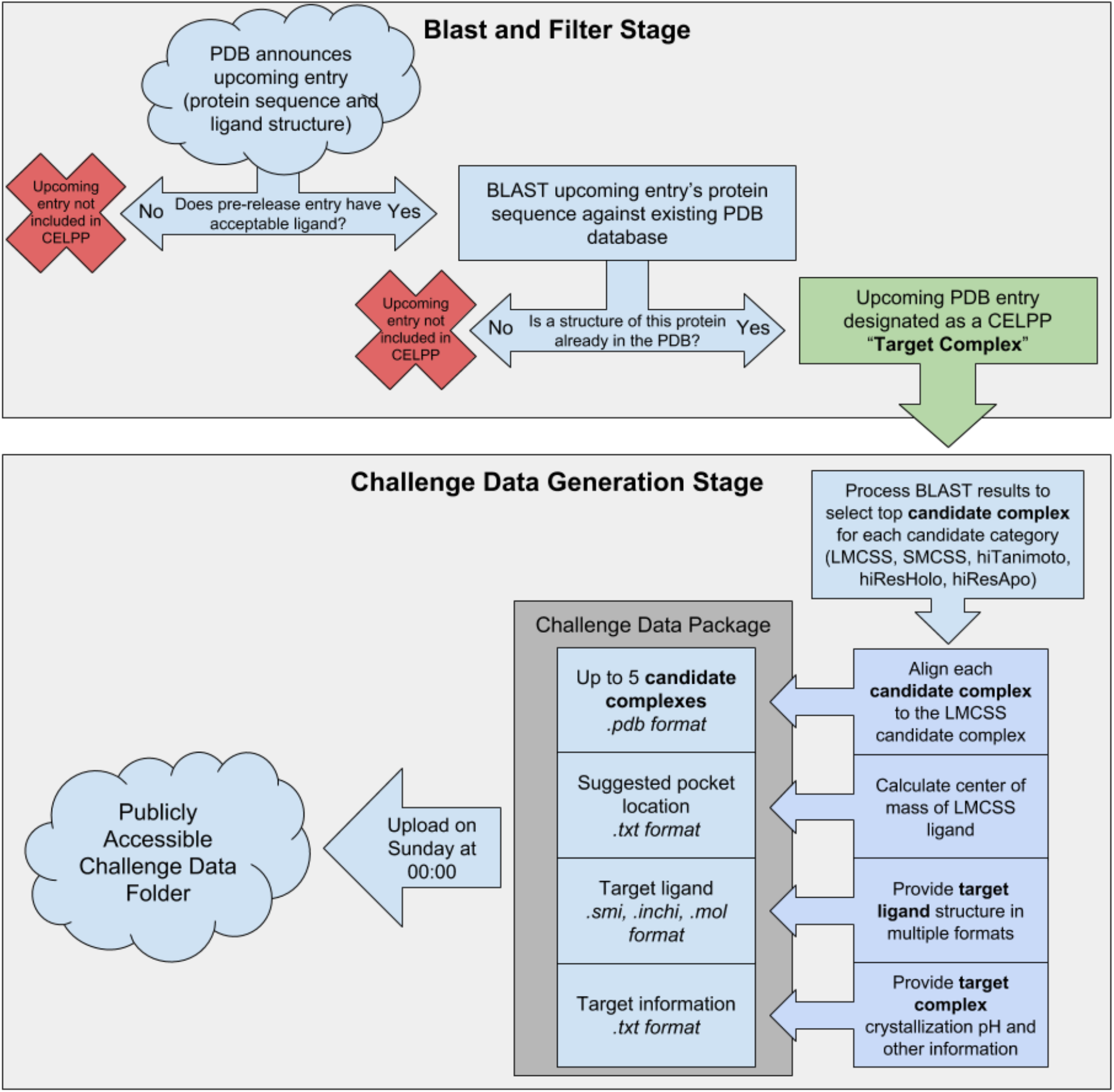
CELPP Target Selection and Challenge Package Generation. CELPP downloads the publicly-available PDB pre-release information and then processes the new entries to assemble the weekly challenge package. Boxes and arrows indicate processing steps, two-way arrows indicate filtering steps, clouds indicate internet-accessible file platforms, and the dark grey box indicates the weekly challenge data package, in which each target is one subdirectory. See main text for details.

Candidate structures for use in the docking challenge are identified by using the sequence comparison program blastp^96^ to find PDB entries with >95% sequence identity and >90% sequence coverage of the target sequence (Figure 2). The resulting proteins are then further filtered to a set that were determined by X-ray crystallography (rather than NMR, for example) and which comprise only a single unique protein sequence. For each target, up to five candidate structures meeting these criteria are selected from the PDB for use as receptors in the cross-docking challenge. The five candidates, which are chosen to test the effects of various criteria for selecting the receptor used in cross-docking, are as follows:

- **Largest Maximum Common Substructure (LMCSS):** This candidate complex contains the ligand with the largest maximal common substructure to the target ligand. The center of mass of the ligand in this complex is used to suggest the binding pocket for all five candidates for this target. In the case that two candidate complexes tie for the largest maximal common substructure, the highest-resolution candidate structure is used.
- **Smallest Maximum Common Substructure (SMCSS):** This candidate complex contains the ligand with the smallest maximal common substructure to the target ligand. In the case that two candidate complexes tie for the smallest maximal common substructure, the highest-resolution candidate is used.
- **Highest Tanimoto Similarity (hiTanimoto):** This candidate complex contains the ligand with the highest ligand Tanimoto score, using the RDKit default fingerprint method,^97^ to the target ligand. In the case that two candidate complexes tie for the highest mutual Tanimoto score, the highest-resolution candidate is used. This candidate sometimes is the same as the LMCSS candidate.
- **Highest Resolution Holo (hiResHolo):** This candidate complex has the highest crystallographic resolution limit of any determined with a druglike ligand.
- **Highest Resolution Apo (hiResApo):** This candidate receptor structure has the highest crystallographic resolution limit of any for this protein determined with no druglike ligand.

#### 3.2.2 Generation of the challenge data package

After the processing described in Section 3.2.1 has been completed, the results are incorporated into a common data package for use by CELPP participants. This typically becomes available in a public Box.com folder by 00:00 Pacific Time on Sunday (Figure 1). For each target, the challenge data package (Figure 2) contains:

- Structures of the candidate proteins in PDB format, aligned to the LMCSS structure coordinates but otherwise unmodified from the PDB entries from which they were drawn
- PDB structure of the LMCSS ligand, drawn from the LMCSS candidate structure
- The suggested binding pocket center (center of mass of the LMCSS ligand) in .txt format
- SMILES, InChI, and 2D MOL files of the target ligand
- A parseable text file containing the PDB ID of the forthcoming entry, the crystallization pH, the HETID of the target ligand, and additional information about the candidate cocrystal structures, such as their PDB IDs and crystallographic resolution limits. See Scheme S1 for sample.

#### 3.2.3 Evaluation of Predictions

After the close of the submission window and release of the experimental cocrystal structure by the PDB (Figure 1), automated scripts evaluate the pose predictions by calculating the symmetry-corrected RMSD of each predicted ligand pose relative to the crystallographic pose, using Schrödinger and OpenEye tools (Scheme S2). When the crystal structure has multiple instances (protein chains) of the target protein, the predicted pose is assigned the lowest RMSD that can be achieved by aligning the predicted protein-ligand complex to each instance of the chain, as detailed in Scheme S2.

### 3.3 Participating in CELPP

#### 3.3.1 Enrollment and Information

In order to obtain upload/download credentials for CELPP data, one must register as a CELPP participant. Information for how to participate in CELPP is available at the D3R website (https://drugdesigndata.org/about/celpp), including links to a CELPP Developers User Group on Google Groups and to the CELPP GitHub Wiki.

#### 3.3.2 Developing a Prediction Workflow

Based on the typical throughput of CELPP challenges, CELPP participants should construct a pose-prediction workflow that can process up to 100 targets in the 63-hour submission window (Figure 4A). This requires executing up to 100 ligand preparation tasks, 400-500 protein preparation tasks, and 400-500 docking tasks. Although CELPP participants may choose to submit results for only a subset of the targets posed each week, doing so makes it difficult to compare different methods on an equal footing, so participants are encouraged to work with all targets each week. To help participants construct their workflows, D3R provides a workflow template, called CELPPade (Figure 3, https://github.com/drugdata/cookiecutter-pycustomdock). CELPPade handles the download, unpackaging, repackaging and upload of the CELPP challenge data, letting the participant focus on implementing their docking solution by writing a few specific methods in Python.

**Figure 3.**
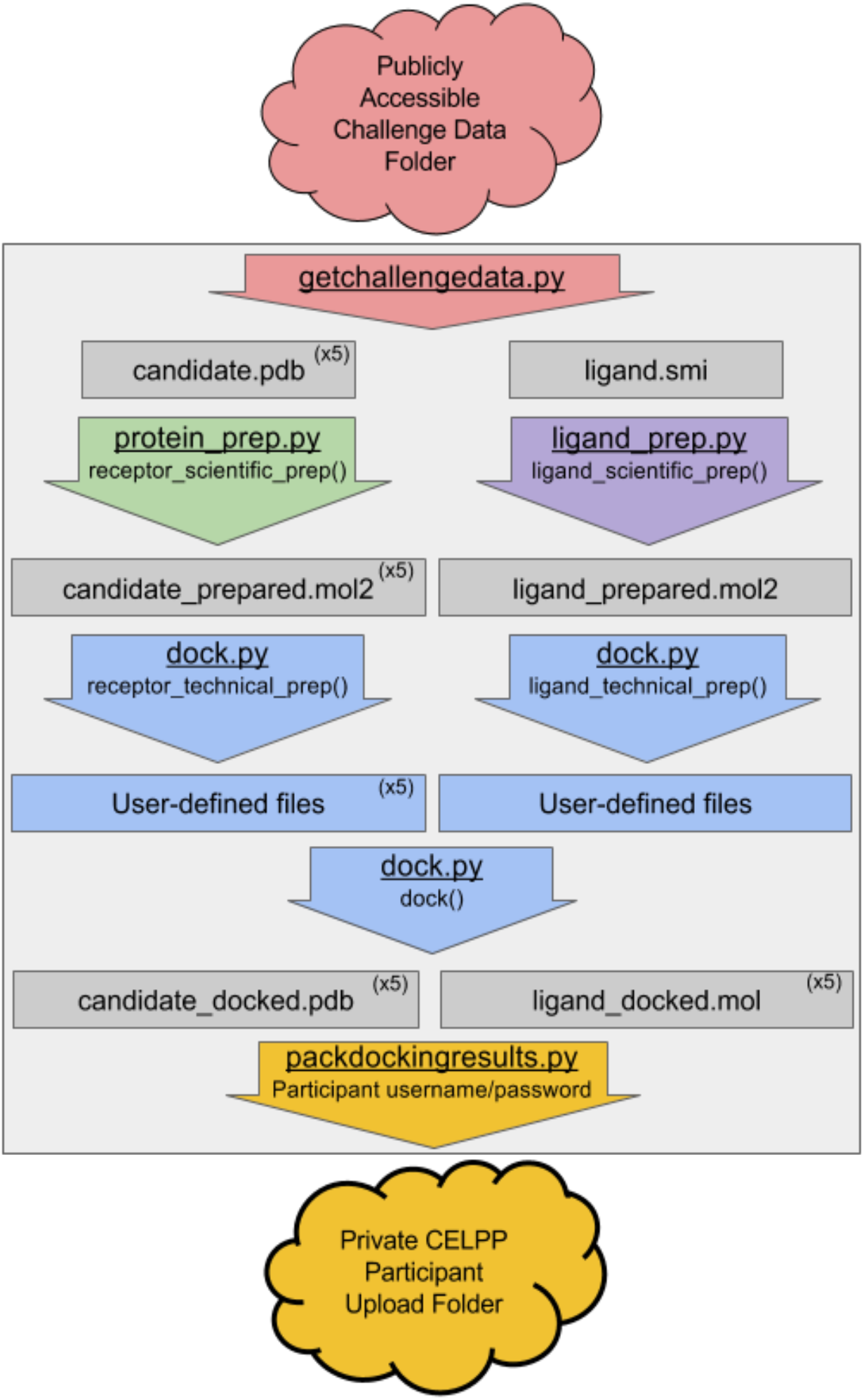
The CELPPade workflow template. Vertical arrows indicate functions, rectangles indicate files passed between steps, and clouds represent internet-accessible folders. The large grey box indicates the steps that are run on the participant’s computer. Different colors indicate script files for different steps of pose prediction, and names ending in () indicate python functions that are implemented by participants. The output files from protein_prep.py and ligand_prep.py is not strictly required to be in mol2 format, but adopting this format will improve interoperability of steps from diverse workflows.

The five user-written Python functions in CELPPade are located in three files (Figure 3, protein_prep.py, ligand_prep.py, and dock.py). These five functions mirror the stages in a general pose prediction workflow: protein and ligand structure preparation, protein and ligand file format conversion, and docking. Each function receives input filenames as arguments, and participants are responsible for populating the function bodies with commands to run the respective stage of their workflow. Participants can implement their docking workflow in these functions using Python commands, or using Python’s interfaces to the command line to execute shell commands. Once implemented, CELPPade is able to run these functions in sequence on each prediction target in the current CELPP challenge week. Combined with the download and upload scripts bundled in CELPPade, implementing these five functions results in a functioning CELPP workflow.

We also provide a tutorial that follows the creation of a model docking workflow based on CELPPade. This docking workflow uses Chimera DockPrep^98^ to prepare both the protein and ligand, and AutoDock Vina to carry out pose prediction.^20, 99, 100^ The model workflow provides examples of running shell commands from within Python, uses software that is free for use by academic labs, and can run on most 64-bit Linux systems with Python, Chimera, RDKit, OpenBabel, and Autodock Vina installed.^98–101^ Note that participants are not required to use the CELPPade template; it is provided only as a convenience.

Even if a participant does not use the full CELPPade package, two helper scripts it contains may be of interest. The first, **`getchallengedata.py`**, helps participants access the correct challenge data package each week. It reads the Box.com login credentials of participants from the user’s customized file `ftp_config` and uses these to access the online folder. It then reads the file `latest.txt` file in the Box.com folder to determine the name of the most recent challenge package, downloads the package, and unzips it in the user’s local folder. The last script, **`packdockingresults.py`**, takes as input a formatted directory of docking results generated by the participant, compresses the directory into a tar file, and uploads the tar file to the participant’s private submission folder. These scripts facilitate automation by providing straightforward upload/download functionality for CELPP data, independent of the specific details of the prediction workflow.

#### 3.3.3 Definition and Submission of Predictions

For each pose prediction, for a given candidate structure, a valid submission comprises the receptor structure in PDB format and the ligand structure in MOL format,^102, 103^ with ligand coordinates in the receptor frame of reference. Participants may choose not to use any of the provided candidate receptor structures for docking but must inform us in this case. Strict adherence to these file formats is required, and deviation from them may result in improper scoring or disqualification of the submitted prediction. Using the CELPPade workflow template will ensure that the docking results directory is appropriately formatted for upload. Pose predictions are uploaded to an online, password-protected Box.com folder provided by D3R. This upload must be completed before 15:00 U.S. Pacific time on Tuesday to be considered valid for scoring.

#### 3.3.4 Score Reporting

Scores are emailed directly to participants and, in the near future, will be publicly accessible at the CELPP website. Participants may choose to remain anonymous, in which case their methods and results will be posted without identifying information.

**Table 1.** Baseline docking workflows. Methods used for protein preparation, ligand preparation, and docking in the D3R inhouse workflows. (Versions: Chimera 1.10.1, RDKit 2016.3.3, MGLTools 1.5.7, AutoDock Vina 1.1.2, Schrodinger 2015-3 release, Omega 2.5.1.4, FRED 3.0.1, RBDock 2013.1/901)

| Workflow Name | Protein Prep Method | Ligand Prep Method | Docking Algorithm |
| --- | --- | --- | --- |
| **Autodock Vina** | Chimera DockPrep<sup>98, 99</sup> and AutoDock Tools prepare_receptor4.py<sup>104</sup> | RDKit 3D coordinate generation,<sup>97</sup> Chimera DockPrep,<sup>98, 99</sup> and AutoDock Tools prepare_ligand4.py<sup>104</sup> | AutoDock Vina<sup>20</sup> |
| **Glide** | Schrödinger PrepWizard<sup>88, 105</sup> | Schrödinger LigPrep<sup>106</sup> | Schrödinger Glide SP<sup>107-109</sup> |
| **OE Fred** | HETATM removal and OpenEye receptor_setup<sup>110</sup> | OpenEye Omega<sup>111</sup> | FRED<sup>68, 110, 112</sup> |
| **rDock** | HETATM removal, Chimera DockPrep,<sup>98, 99</sup> rbcavity<sup>113</sup> | RDKit 3D coordinate generation<sup>97</sup> | rbdock<sup>113</sup> |

### 3.4 Baseline docking workflows

To illustrate CELPP, test our procedures, and provide a baseline of performance, we have used the CELPPade template (Section 3.3.2) to create four in-house CELPP workflows, based on the Autodock Vina,^20^ FRED,^68, 112^ Glide,^107, 108^ and rDock^113, 114^ docking suites. These use the protein and ligand preparation tools that accompany the respective docking codes where possible, and open-source tools otherwise (Table 1). These workflows represent default implementations of their respective methods. To standardize the testing, all workflows are set to consider the same docking region. Algorithms that require a docking box are set to use a 15×15×15 Å region, and software that requires a sphere is set to use a 10 Å radius region. As we have made no effort to optimize the workflows, their performance may not be reflective of the best performance the algorithms can provide. All of the in-house prediction workflows are available on GitHub. In addition, the AutoDock Vina workflow has been documented in a tutorial on CELPP workflow development. This can be accessed on the D3R GitHub page, as noted in Section 3.3.2.

## 4 Results

### 4.1 Scale and Character of the CELPP Challenge

During a 66-week period spanning parts of 2017 and 2018, 1,989 targets met the CELPP criteria (Section 3.2.1) and were submitted to outside participants and our in-house workflows (Section 3.4). To permit meaningful analysis, the initial data discussed in this paper have been filtered to include only targets for which at least 3 workflows submitted predictions in the LMCSS category, and at least one LMCSS prediction achieved an RMSD under 8 Å (Table S1). Future analyses will allow more sophisticated selections and comparisons. Most weeks saw 20-50 targets, and the maximum number of targets per week has remained below 100 (Figure 4, top). The ligands to be docked had an average of 27 heavy atoms and 5 rotatable bonds. The distributions of these descriptors are provided in the bottom panel of Figure 4, and the full list of PDB IDs and ligand SMILES strings is provided in the SI. In some cases, the selection criteria for the candidate categories yielded the same PDB structure in different categories for a target complex. For example, the PDB structure with the largest maximum common substructure (LMCSS) to the target ligand may also be the one with the highest Tanimoto similarity index (hiTanimoto). The frequency of these overlaps is shown in Figure S1. Because apo structures are not always available, there are fewer candidate structures in that category.

**Figure 4:**
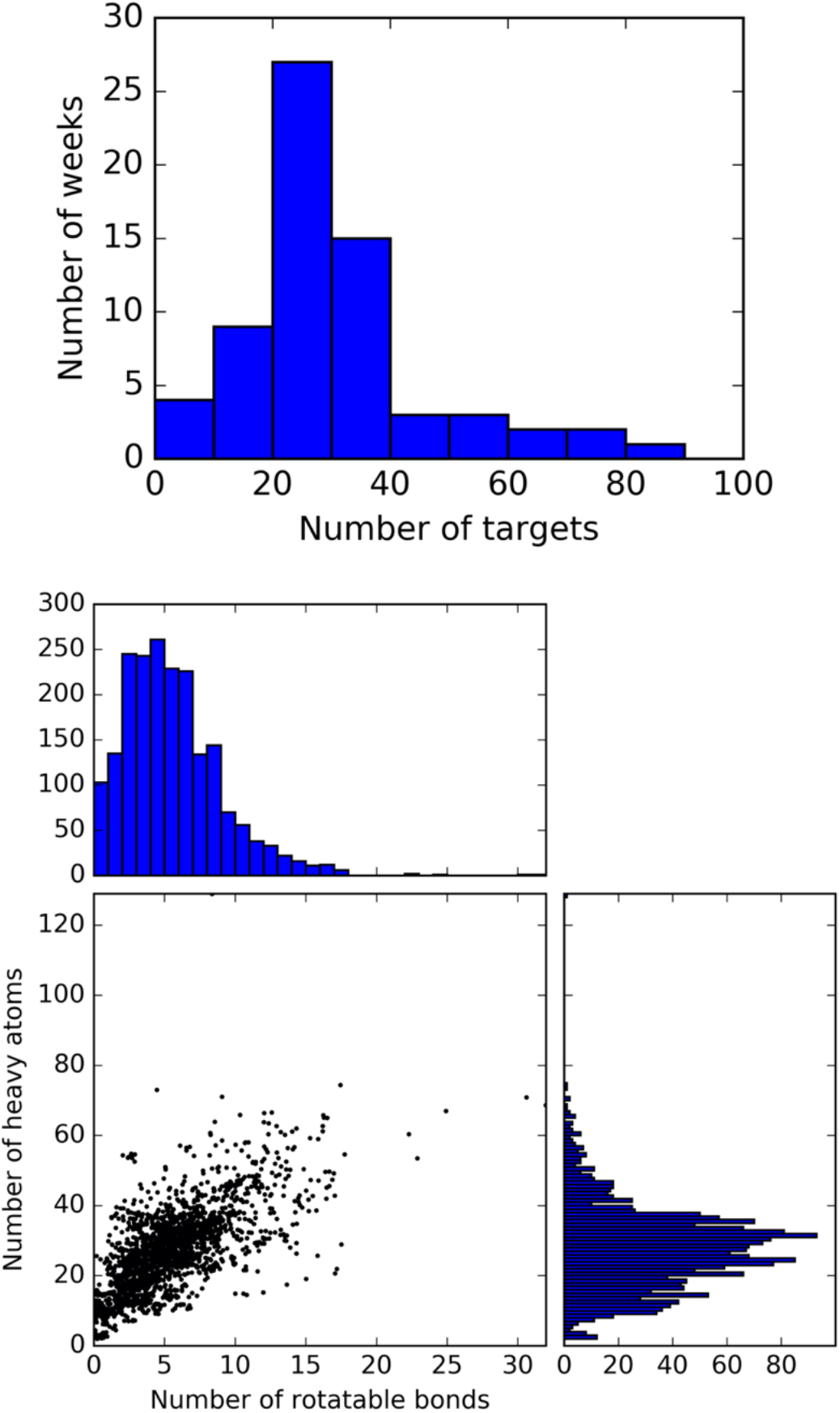
Characteristics of CELPP weekly challenges. Top) Number of CELPP targets per week (66 weeks total). Bottom) Characteristics of CELPP target ligands (n=1,989). Each dot represents one target ligand, and histograms above and to the right show the distribution of characteristics on each axis. Uniformly distributed random values in the range [−0.5, 0.5] were added to X and Y coordinates to better show point density. Numbers of rotatable bonds and heavy atoms were calculated from InChI strings using RDKit.^97^

### 4.2 Pose Prediction Performance to Date

The performance records of three anonymous early-adopting external participants (one of them submitting the results from two distinct workflows) and the four in-house workflows over a 66-week period spanning 2017 and 2018 already provide number of informative and illustrative analyses. Previous studies of pose prediction have generally considered a ligand RMSD within 2 Å of the crystal pose to be useful for compound design.^20, 45–47^ In the CELPP dataset, the median prediction RMSD for the best-case prediction categories (LMCSS and hiTanimoto) is around 5 Å (Figure 5, middle). In these categories at best 20% of pose predictions are accurate to within 2 Å RMSD, and only about 40% are accurate to within 4 Å RMSD (Figure S2). These rates are significantly worse than those seen in a prior blinded challenge,^47^ where about 34% of the top scoring poses generated by various docking codes had RMSDs less than 2 Å, when averaged across all protein targets. However, it is important to note that all participants in the prior study were provided with receptor structures hand-picked and prepared by human experts to accommodate the ligands to be docked. In contrast, CELPP receptors are selected automatically and are not prepared by system experts. The CELPP success rates are on par with those found in the pose-prediction components of the recent D3R Grand Challenge 3, which yielded a corresponding success rate of 16%. Much as in CELPP, Grand Challenge participants are not provided with expertly selected and prepared receptor structures.

**Figure 5.**
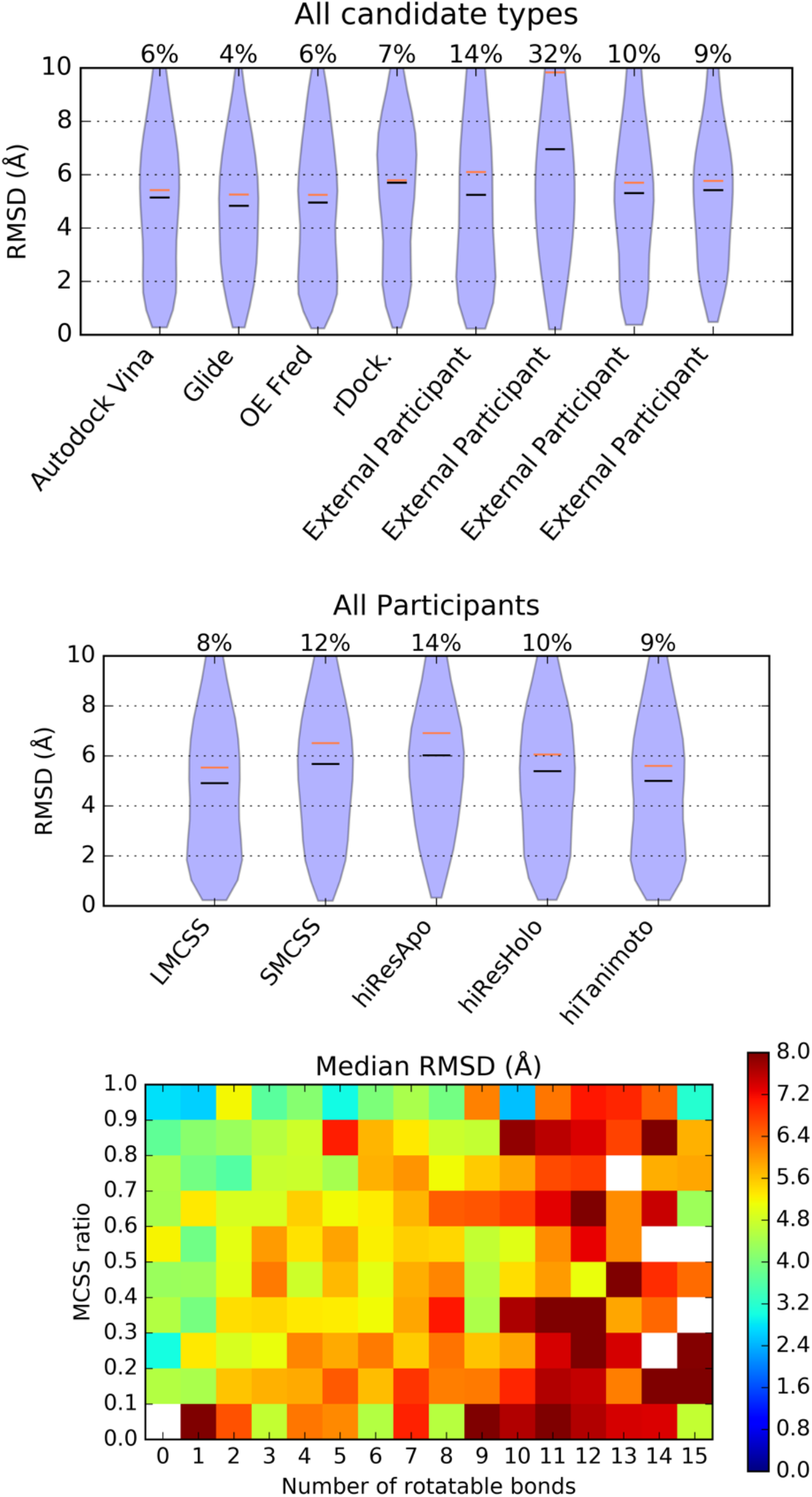
Results of CELPP. Top) Performance by participant or in-house method, combining predictions from all candidate categories. Middle) Performance by candidate category, combining predictions from all participants and in-house methods. Black line indicates median, orange line indicates mean. The number above each plot indicates the fraction of predictions above 10 Å. Bottom) Median prediction RMSD as a function of number of rotatable bonds and MCSS ratio. The MCSS ratio is defined as the fraction of the heavy atoms in the target ligand that are in its maximal common substructure with the candidate ligand. Data are taken from all participant and in-house method predictions in LMCSS and SMCSS categories. White indicates no data.

A finer-grained analysis reveals that most methods provide rather similar levels of accuracy (Figure 5, top), based on median RMSD, with rDock and one External Participant performing noticeably worse. Focusing on the performance of various methods when provided high-similarity structures (LMCSS and hiTanimoto categories; see next paragraph), we observe that two anonymous public participants performed slightly better than other methods, as measured by the fraction of RMSDs in the 0-2 Å range (Figure S2). The in-house OE Fred and Vina workflows yield rather similar results, with GLIDE variants and rDock trailing slightly by this metric. As noted above, the in-house workflows are not tuned for optimum results and thus may not reflect the best performance these algorithms can provide.

The extensive data set provided by CELPP also allows quantification of important trends that have been previously noted.^45, 46^ First, docking into a receptor determined with a chemically similar ligand, as determined by the maximum common substructure (LMCSS) or the fingerprint Tanimoto similarity index (hiTanimoto), more than doubles the success rate (RMSD < 2Å), relative to docking into a structure determined without a bound ligand (hiResApo) (Figure S2). Docking into the highest resolution structure solved with any ligand (hiResHolo) and into the structure with the least similar ligand, based on maximum common substructure (SMCSS), yielded results of intermediate accuracy. Similar results are observed for the individual methods (Figure S2). Second, docking results tend to be less accurate for ligands with more rotatable bonds, but this challenge can be overcome by docking into a protein structure determined with a highly similar ligand (Figure 5, bottom). Best docking results are, therefore, obtained either when the number of rotatable bonds is less than 2 or when the fraction of the heavy atoms in the target ligand that are in its maximal common substructure with the candidate ligand (the MCSS ratio) is above 0.8. The worst results are obtained for target ligands with >10 rotatable bonds and an MCSS ratio lower than about 0.5. In the best-case scenarios, where the candidate structure has 0 or 1 rotatable bonds and an MCSS ratio of at least 0.8, automated docking workflows can achieve a median RMSD of around 3 Å. In more difficult cases, with MCSS ratios between 0.4 and 0.5 and 7 rotatable bonds, the median RMSD rises to about 6 Å.

The “standard” target, 1FCZ, is included in the challenge package each week. As 1FCZ is an existing PDB structure, the LMCSS category poses a self-docking challenge where the correct ligand pose is publicly known (results from 1FCZ are excluded from the general dataset). All workflows regularly achieve RMSD values below 1 Å in the LMCSS category for 1FCZ (Figure S3). Three internal and two external workflows behave deterministically on this target, returning the same poses each week. One internal workflow (rDock) and two external workflows return different poses each week. These inconsistent results indicate a potential source of uncertainty for method comparison. The Glide workflow implemented in CELPP does not produce a prediction for 1FCZ, as the size of the docking region used for all methods in this study is smaller than its recommended value. However, when run with its default size docking box, the GLIDE workflow consistently produces a pose with an RMSD of 0.4 Å.

## 5 Discussion

The CELPP challenge introduced here is a powerful new cyberinfrastructure tool to evaluate and improve protein-ligand pose prediction technologies. Unlike prior blinded prediction challenges in this field, CELPP sets a new challenge each week, each with dozens of new ligand-protein complexes to model, and provides rapid and consistent feedback for participants. The >1,900 individual challenge cases set by CELPP in one 66-week period far exceeds the number of cases set by all prior blind pose-prediction challenges, and the CELPP challenge is ongoing.

The high throughput nature of CELPP provides a dramatic increase in the statistical power of analyses for docking, and thus enables sharper distinctions among methods. We anticipate new insights into not only the core docking algorithms, but also key procedural details, such as how crystallographic water molecules and protonation states are treated. We also plan to look for characteristics of protein targets and ligands that correlate with the performance of specific methods. For example, some algorithms may do better for hydrophobic sites or for specific protein families, such as serine proteases. Such statistical analyses will help practitioners choose methods suited for their specific applications and set meaningful expectations for the quality of predictions on new systems. In parallel, we will scan for cases where multiple methods do poorly, checking for situations in which CELPP’s automated procedures may generate inappropriate challenges, such as where a cofactor is present in the candidate but not in the target. The volume and tempo of the CELPP challenge also allow its use in iterative optimization of pose prediction methods. Thus, we anticipate that CELPP will help drive the development of increasingly predictive docking workflows.

It is worth noting that, if a participant’s pose-prediction method were to change over time, it would become impossible to collect full statistics for a single, defined method. It will therefore be useful to distinguish between those methods which are locked in for a period of time, and for which meaningful statistics therefore can be obtained, from those which are mutable, such as when CELPP is used to guide ongoing improvements in a pose-prediction method. Perhaps the best way to address this will ultimately be for participants to package their stable methods into shareable workflows, using technologies such as Docker^115^ and Singularity^116^, which can then be executed automatically on machines hosted by the CELPP project. Participants also would benefit by not having to manage the processes or allocate computer time for the calculations. Ideally, such workflows would produce consistent results and be structured into modular steps with standardized I/O, as this would enable the creation and benchmarking of new strategies that recombine steps from various workflows. Such derivatization of workflows could, for example, make it possible to evaluate how various ligand-preparation methods affect the quality of the final pose predictions. This direction promises to build a beneficial culture of generating methods that are rigorously evaluated through ongoing blinded challenges and that are readily shared, so that effective methods can easily be put to use.

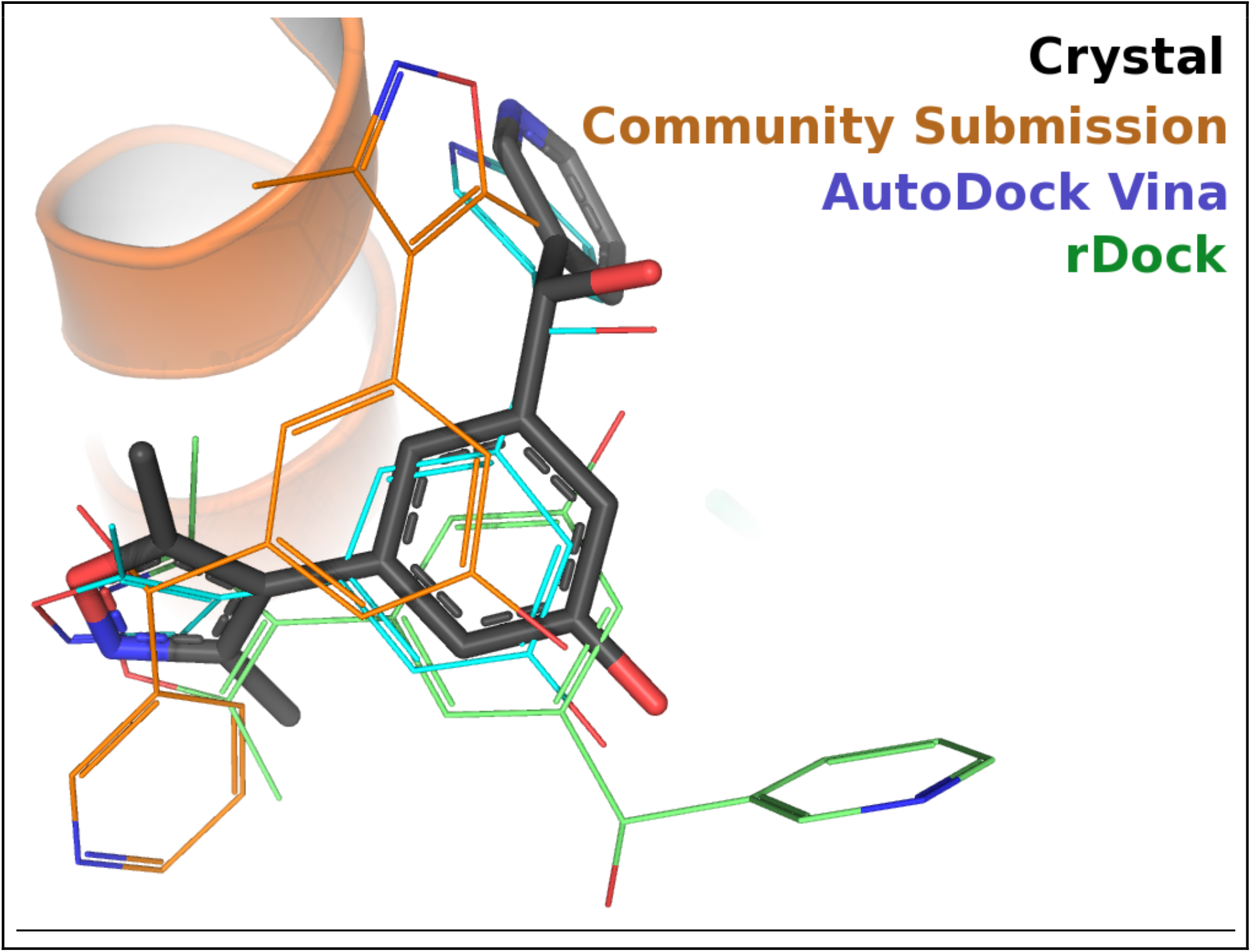

## 6 Acknowledgements

We gratefully acknowledge the early CELPP participants, who provided feedback and developed automated workflows before the challenge was made public. We thank Torsten Schwede and Jürgen Haas for helpful discussions and for developing the inspirational CAMEO challenge. D3R is supported by NIH grant U01 GM111528 to REA and MKG. The contents of this paper are solely the responsibility of the authors and do not necessarily represent the official views of the NIH.

